# Unifying the Ambivalent Mechanisms of Protein-Stabilising Osmolytes

**DOI:** 10.1101/2020.02.22.960856

**Authors:** Mrinmoy Mukherjee, Jagannath Mondal

## Abstract

The mechanism of protein stabilization by zwiterionic osmolytes has remained a long-standing puzzle. While the prevalent mechanistic hypothesis suggests an ‘osmophobic’ model in which osmolytes are assumed to stabilize proteins by preferentially excluding themselves from the protein surface, emerging evidences of preferential binding of popular osmolyte trimethyl amine N-oxide (TMAO) with hydrophobic macromolecules contradict this view. Here we address these contrasting perspectives by investigating the folding mechanism of a set of mini proteins in aqueous solutions of two different osmolytes glycine and TMAO, via free energy simulations. Our results demonstrate that, while both osmolytes are found to stabilize the folded conformation of the mini proteins, their mechanism of actions are mutually diverse: Specifically, glycine always depletes from the surface of all mini proteins, thereby conforming to the osmophobic model; but TMAO is found to display ambivalent signatures of protein-specific preferential binding and exclusion to/from the protein surface. At molecular level, the presence of an extended hydrophobic patch in protein topology is found to be recurrent motif in proteins leading to favorable binding with TMAO. Finally, an analysis based upon the preferential interaction theory and folding free energetics reveals that irrespective of preferential binding vs exclusion of osmolytes, it is the *relative* preferential depletion of osmolytes on transition from folded to unfolded conformation of proteins, which drives the overall conformational equilibrium towards the folded state in presence of osmolytes. Taken together, moving beyond the model system and hypothesis, this work brings out ambivalent mechanism of osmolytes on proteins and provides an unifying justification.

## Introduction

Osmolytes are small cosolutes which protect the protein’s native tertiary fold in adverse denaturing condition and support cells to cope up with the osmotic stress.^1–5^ Over the years, several zwitterionic osmolytes, namely Trimethyl amine N-oxide (TMAO), glycine, glycine betaine have emerged as efficient stabilizers of protein. One of the most popular models at the forefront of hypothesizing the mechanism of action of protein-stabilizing osmolytes is the so-called ‘osmophobic’ model. Originally proposed by Bolen and coworkers,^6,7^ the model bases its hypothesis upon free energy change of amino-acids of proteins on their transfer from neat water to osmolyte solution. This model had suggested that osmolytes’ protein-stabilizing ability stems from a delicate balance of their unfavorable interaction with the protein backbone and favorable interaction with the side chains, with the backbone contributions dominating the side chain contributions. In this mechanism, all zwitterionic osmolytes are considered to be depleted from the protein surface and generally assumed to conform to this osmophobic model. Over the years, this model of preferential exclusion of osmolytes from protein surface has received extensive supports in multiple investigations. ^8–14^ However, some recent investigations^15–18^ involving the quintessential osmolyte TMAO are beginning to hint at its more complex and heterogenous molecular picture of the protein-stabilizing role, showing signs of deviation from the tenets of popular osmophobic model. The present work provides convincing evidence of the ambivalent mechanism of osmolytes’ stabilizing action towards a set of multiple mini proteins and subsequently unifies these observations within the frame-work of preferential interaction theory.

In one of the earliest works signaling the deviation of osmolytes’ mechanism of action from the usual preferential exclusion based osmophobic model, Mondal et al.^15^ simulated the collapse behavior of a model hydrophobic polymer and had proposed the idea of TMAO’s stabilization of collapsed ensemble of hydrophobic polymer via the favorable binding of TMAO to the polymer surface, as opposed to the usual hypothesis of osmolyte’s preferential exclusion from the macromolecular surface. Via a combination of computer simulation of collapse behavior of a model bead-on-a-spring hydrophobic polymer and Wyman-Tanford preferential binding theory,^19,20^ the work had interpreted that TMAO more favorably binds to the collapsed conformation of the hydrophobic polymer surface than the extended conformation. This work analyzed the preferential binding coefficient of the osmolyte with the model polymer and based its argument in terms of the Wyman-Tanford preferential binding theory.^19,20^ It showed that what matters is the higher extent of preferential binding of TMAO with the collapsed conformation compared to that with an extended conformation, which guides the direction of conformational equilibrium of a hydrophobic polymer towards the collapsed state. This hypothesis of stabilization of hydrophobic interaction by osmolytes via its preferential binding has received subsequent experimental validation ^21^ on osmolyte-induced collapse of synthetic polymer polystyrene. Since then, the idea of stabilization of hydrophobic interaction via preferential binding of TMAO with the surface, in contrast to its exclusion from the surface, has received regular attention in multiple subsequent studies.^18,22–29^ However, these works have been mostly limited to model polymer and not been validated in case of realistic proteins, until recently.

A recent report of osmolytes’ action towards hydrophobic protein elastin^16^ had signalled a departure of protein-osmolyte interaction from the osmophobic model. Using lower critical solution temperature measurements, surface tension measurements and Infra-red spectroscopy, this work showed evidence of TMAO’s favorable binding with the protein surface, in contrast to the preferential exclusion of glycine and glycine betaine from the same peptide. However, in lieu of any direct evidence of the *conformation dependence* in the preferential interaction of osmolytes with the protein surface, this work referred to earlier report and analysis on model polymer^15^ as a possible mechanism at play in protein and mainly speculated on the possibility of conformation dependence in preferential exclusion or binding of these osmolytes in folded versus unfolded conformation of the protein. On the other extreme, very recent effort^18^ by current authors of the present manuscript, albeit on model hydrophobic polymer, reinforced Cremer and coworkers’^16^ observation that osmolytes’ mechanistic action towards hydrophobic interaction can be quite heterogeneous: while glycine was found to conform to the usual rule of preferential exclusion from the polymer surface, TMAO was found to show interesting trend of preferential binding with the polymer surface. The work further went on to identify the collapsed and extended ensembles of the hydrophobic polymer and explicitly showed that the extent of preferential interaction (irrespective of exclusion or binding) is conformation-dependent.

Taken together, the recent studies either have completely focussed on model polymer system or have relied on the analysis of model system to interpret the observations in typically hydrophobic protein. These keep open a pertinent question: Is it possible to establish a protein-specific structural motif to explain the ambivalent mechanism of osmolytes towards proteins and to explore the conformation-dependent preferential interaction of the osmolytes with the protein? The current work closes this gap by computationally simulating the full folding landscape of a set of popular mini proteins in two different osmolyte solutions, namely glycine and TMAO. We report that while both the aqueous solution of osmolytes stabilize the folded ensemble of the mini proteins, the mechanism of action of glycine and TMAO towards the mini proteins can be different. Specifically, we find that glycine always depletes from the surface of all mini proteins, thereby conforming to the osmophobic model. However, on the other hand, we find that TMAO can display the ambivalent nature of possible binding and depletion from the surface of the side chains of the mini proteins. Mapping the side chain preferential interaction coefficients with the hydrophobicity index of the side chain residues shows a strong correlation of preferential binding of methyl groups of TMAO with the hydrophobic residues. More importantly, the presence of an extended patch of hydrophobic residues is found to be a generic motif for mini proteins which can lead to favorable binding with TMAO. Finally, we individually analyze the preferential interaction profile of TMAO and glycine with both folded and unfolded ensembles and reveal that irrespective of preferential binding vs exclusion of osmolytes, it is the relative difference in the preferential binding/exclusion coefficient of osmolytes with/from the unfolded and folded ensembles, which drives the mechanism of protein stabilization in both osmolytes. Taken together, moving beyond the model system and hypothesis, this work brings out an ambivalent behavior of osmolytes with real proteins and unifies the heterogenous mechanisms of osmolytes within the framework of preferential interaction theory.

## Results and Discussion

### Both Glycine and TMAO stabilise folded ensembles of small proteins

Trimethyl amine N-oxide (TMAO, Figure 1H) and glycine (Figure 1G) have long been perceived as the classic protein stabilizers. As a proof of concept, we first investigate the effect of aqueous TMAO solution on the biomolecular conformational landscapes. Towards this end, we compare the free energy landscapes of two mini proteins, namely GB1 (Figure 1A) and Trp-cage (Figure 1B) in neat water, in aqueous TMAO solution and in aqueous glycine solution (see Figure 2). A judicious combination of adaptively sampled MD trajectories (spawned by a priori run replica exchange Molecular Dynamics (REMD) simulated conformations) and Markov State Model (MSM) (see Methods) enables us to quantify the free energy landscape of these two proteins in neat water and in aqueous solution of osmolytes. Figure 2A and B compare two-dimensional free energy profiles of GB1 *β*-hairpin (PDB: 1GB1, residues 41 − 56; we call it GB1)^30^ along its radius of gyration (*r*_*g*_) and fraction of native contacts (*NC*) in neat water and in aqueous TMAO solution. Similar profiles for Trp-cage (PDB: 1L2Y)^31^ are shown in Figure 2D and E. Typically a decrease in the value of *NC* of the proteins and an increase in *r*_*g*_ from the folded protein conformation reflect protein unfolding. The choice of *r*_*g*_ and *NC* as the reaction coordinates for free energy landscape stemmed from our previous analysis of optimized reaction coordinates and earlier precedences on their usage in the investigation of protein free energy landscape.^32–34^ Interestingly, as a generic feature of all these free energy profiles computed in this work, we find that the major conformational changes of the proteins are mostly captured along *NC*, while the changes in the *r*_*g*_ of the proteins are secondary. Quite significantly, we find that the unfolded ensembles of both GB1 and Trp-cage become free energetically more unfavorable in aqueous TMAO solution, when compared with that in neat water. The aqueous solution of glycine imparts a qualitatively very similar stabilizing effect (Figure 2C and F), as is observed in aqueous TMAO solution, for both the systems of our interest: Populations of unfolded conformations of both GB1 and Trp-cage get reduced in both osmolyte solution. The overall analysis re-affirms glycine and TMAO’s stabilizing role of the folded ensemble, relative to the unfolded ensemble, in both these small proteins. Hence both these systems potentially would serve as key benchmarks in understanding the molecular mechanism behind these two osmolytes’ role in protein stabilization.

**Figure 1:**
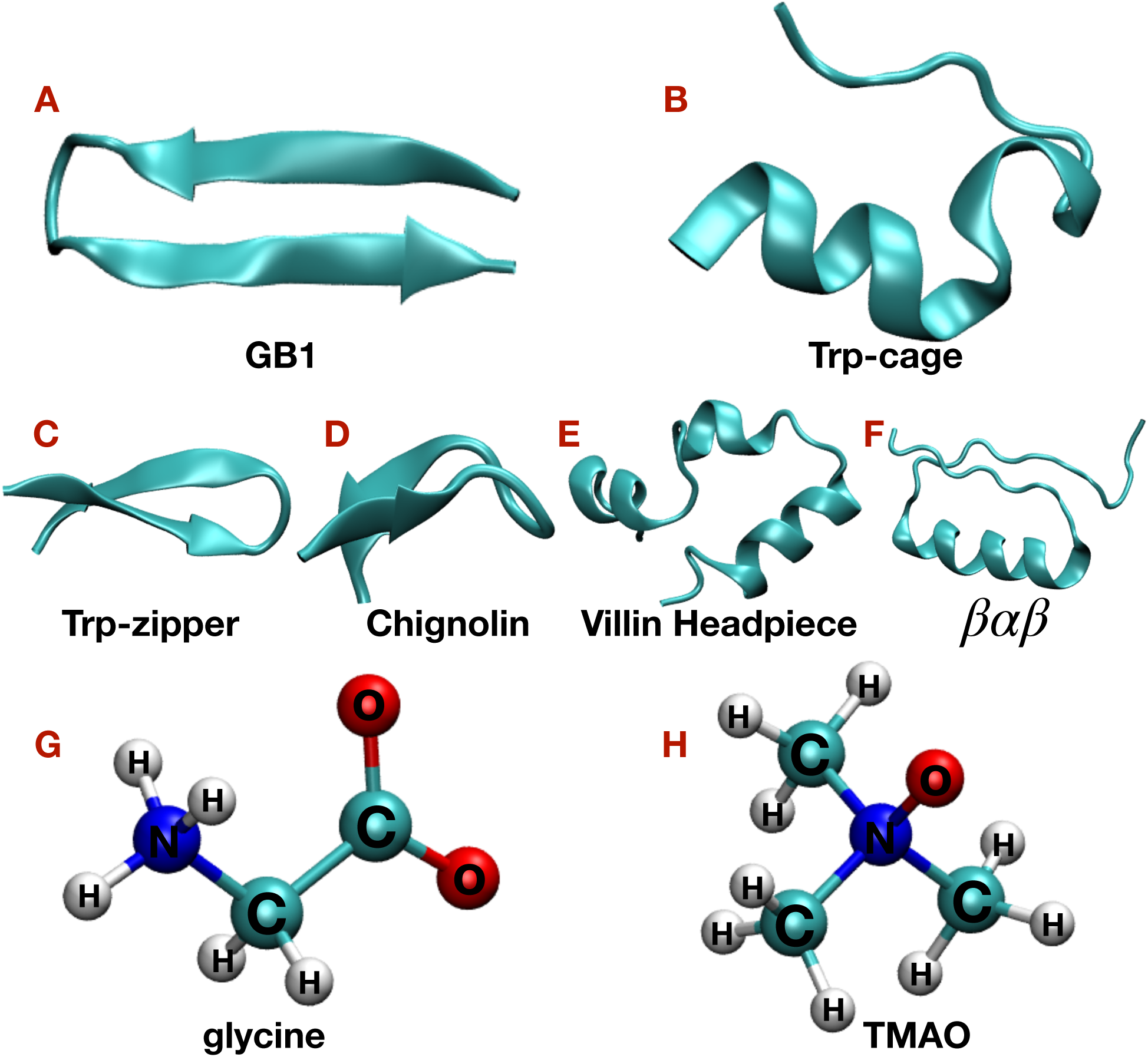
Representative snapshots of native structure of GB1 *β*-hairpin (A), Trp-cage (B), Trp-zipper (C), Chignolin (D), Villin Headpiece (E) and *βαβ* (F) and the structure of osmolyte molecules glycine (G) and TMAO (H).

**Figure 2:**
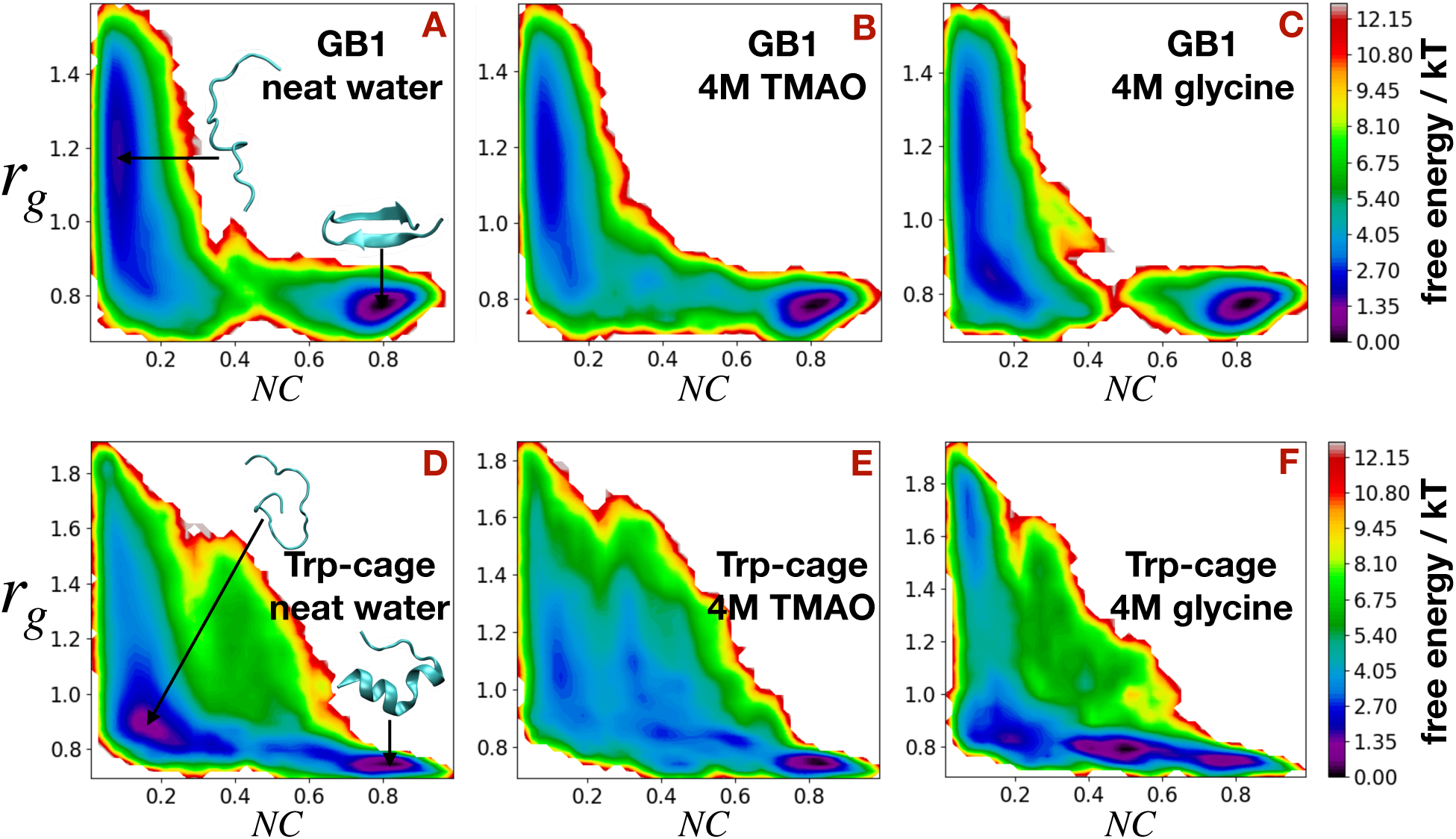
Free energy profiles of GB1 and Trp-cage along its radius of gyration (*r*_*g*_) and fraction of native contacts (*NC*) in neat water (A and D), in aqueous TMAO solution (B and E) and in aqueous glycine solution (C and F). Representative folded and unfolded configurations are shown in the figure.

### TMAO deviates from canonical view of preferential interaction with protein

The canonical view of osmolyte’s action on proteins suggests that the zwitterionic osmolytes are supposed to be preferentially excluded from the protein surface. To investigate this popular hypothesis, we analyze both the osmolytes’ preferential interaction (relative to water) with GB1 and Trp-cage. We employ an experimentally relevant quantity called preferential interaction coefficient (Γ)^19,20^

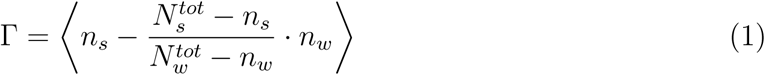

where *n*_*s*_ is the number of cosolutes (glycine or TMAO) bound to the protein and 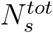 is the total number of cosolutes in the system. On the other hand, *n*_*w*_ is the number of water molecules bound to the protein and 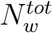 is the total number of water molecules in the system. Γ quantifies the excess of cosolute molecules *s* in the protein solvation shell as compared to its average concentration in the solution. We calculate the profile of Γ as a function of distance (*r*) from the folded conformation of the protein. In these profiles, Γ > 0 indicates the preferential binding of osmolytes to the protein surface, while a negative Γ provides a measure for the depletion of osmolytes from the protein surface. Figure 3A-D compares glycine and TMAO’s distance-dependent preferential interaction profile with both the proteins (GB1 and Trp-cage). As depicted in Figure 3A and B (black curves), glycine displays a negative value of Γ for a wide range of distance (shaded regions in Figure 3) from both GB1 and Trp-cage. In other words, glycine excludes from the surface of both the proteins. This picture of preferential exclusion of glycine from both the protein surface fits very well with the popular hypothesis that protecting osmolytes are supposed to exclude from the protein surface.^7^ Consistent with current results, similar observations of exclusion of glycine from hydrophobic protein elastin^16^ and its derivative glycine betaine from Trp-cage mini protein^35^ have also been previously reported. However, preferential interaction profiles of TMAO with these two systems (Figure 3C and D (black curves)) paint a mutually contrasting picture: Specifically, we find that while TMAO preferentially depletes from the surface of GB1 at all protein-osmolyte separations, its preferential interaction coefficient with Trp-cage becomes significantly positive for distance *r* > 0.55 nm, implying that TMAO preferentially binds to the surface of Trp-cage for a wide range of protein-osmolyte distances. This preferential binding of TMAO with Trp-cage mini protein is an interesting observation and suggests a clear departure from the routine osmophobic model of preferential exclusion hypothesis in osmolytes. We find that the observations of preferential exclusion of TMAO with GB1 and preferential binding of TMAO with Trp-cage are robust with the variation of TMAO force fields (Figure S1 A-B): Across all forcefields tested here, TMAO is found to favorably bind to the Trp-Cage while it excludes from the GB1 surface. We have also varied the force field of Trp-cage between CHARMM36 (currently reported) and AMBER99SB-ILDN.^36^ We find that both forcefields recover the picture of preferential binding of TMAO to Trp-cage surface (Figure S1 C). These observations lend credence to the picture of preferential binding of TMAO with specific protein (and many other proteins as would be shown later in the current article).

**Figure 3:**
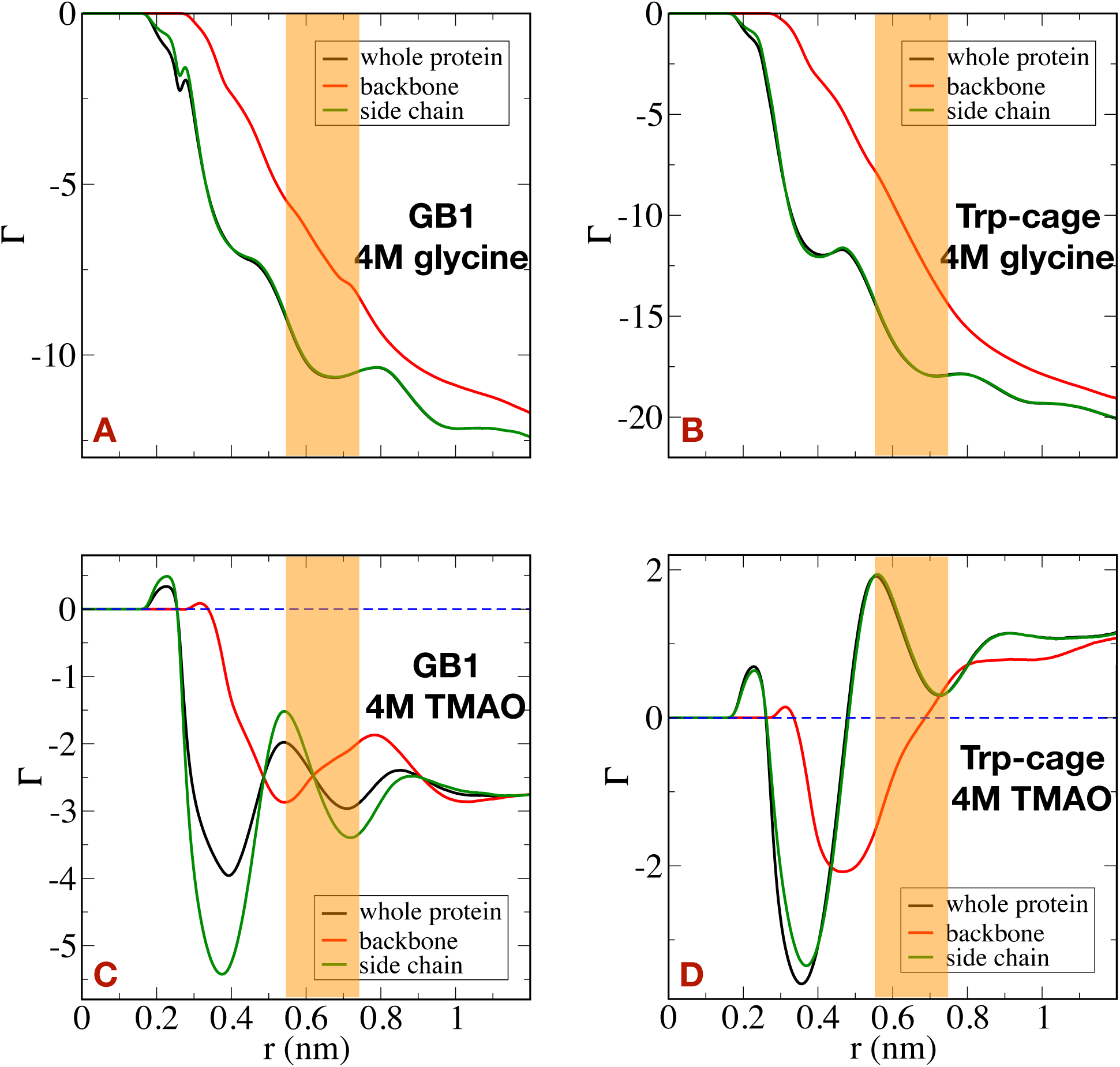
Preferential interaction coefficient (Γ) of glycine (A and B) and TMAO (C and D) with whole part, backbone and side chain of GB1 and Trp-cage as a function of distance (*r*) from the protein surfaces.

To gain further molecular insights into the factors controlling the overall trend of preferential interaction, we dissect the net preferential interaction profiles of both the osmolytes with the whole protein into their respective contributions from backbone and side chain components of the protein. This is relevant as it has been earlier hypothesized by Bolen and coworkers^7^ that it is the unfavorable interaction of zwitterionic osmolytes with the polar backbone of protein surface which sways the balance towards the overall preferential exclusion of these cosolutes from the protein surface. The preferential interaction profiles of whole protein, backbone and side chain are plotted in the same figure.(Figure 3A-D) We find that both TMAO and glycine deplete from the backbone of both GB1 and Trp-cage (Figure 3A-D, red curves), which is in line with the previous hypothesis. However, as an interesting observation of the current work, we also find that the overall trend of preferential interaction profile of both the osmolytes with the whole protein (black curve) is completely dominated by their interaction with side chains of these proteins (Figure 3A-D, green curves). Specifically, we find that glycine excludes from the side-chain of both GB1 and Trp-cage. However, TMAO excludes from the side chain of GB1, while it preferentially binds with the side-chain of Trp-cage. Together, these observations point towards the ambivalent nature of TMAO’s interactions with the side chains of the proteins and necessitates the atomistic exploration of the interplay of side chain diversity with the osmolytes.

### Hydrophobic interaction with TMAO as a key mediator

The current report of TMAO’s preferential binding with the Trp-cage mini protein, especially with the side chain, is reminiscent of recent report^16^ of TMAO’s favorable interaction with the hydrophobic protein Elastin. Mondal et al. have also previously reported TMAO’s favorable interaction with the polystyrene,^21^ hydrophobic polymers. These previous reports on hydrophobic macromolecules and the overall trend of preferential binding of TMAO with Trp-cage and preferential exclusion of TMAO with GB1, as observed in the current work, prompted us to explore the correlation between preferential interaction of side chain of constituent amino-acid residues of GB1 and Trp-cage with the side chain hydrophobicity. Figure 4B and E map the values of preferential interaction coefficient (Γ) of TMAO at *r* = 0.55 nm on the side-chains of amino-acid residues of GB1 and Trp-cage. In the same figure, we mark the side chain of these amino-acid residues according to their hydrophobicity index (Figure 4A and D). The comparison brings out very interesting features: we find that the hydrophobic and aromatic side chains (NP) mostly encourage favorable binding with TMAO, thereby leading to a positive value of Γ. On the other hand, the polar (P) and charged (both electropositive and electronegative) side chains lead to TMAO being preferentially excluded from these side chains. Based on the comparison of the spatial topology of both these systems, we notice an extended patch of hydrophobic residues in Trp-cage, which we believe, plays an important role in its net favorable binding with TMAO. On the other hand, the topology of GB1 indicates an intermittent presence of hydrophobic residues, spatially interspersed by the presence of polar and charged residues, which leads to the net preferential exclusion of TMAO from the surface of GB1. The comparison of TMAO Γ values at *r* = 0.55 nm computed using its constituent atoms (methyl carbon (C), Nitrogen (N) and oxygen (O) atoms) with each amino-acid side chains (Figure 4G) shows that methyl carbon (C) consistently contributes to the most in its preferential binding with the side chain, suggesting TMAO’s propensity for favoring hydrophobic interaction. A similar map of glycine’s preferential interaction with the amino-acid side chains of both GB1 and Trp-cage represents the familiar features of osmolyte’s preferential exclusion with both the proteins. We find that the values of Γ of glycine are mostly zero to negative with most side chains, presumably due to lack of any hydrophobic moieties (Figure 4C and F).

**Figure 4:**
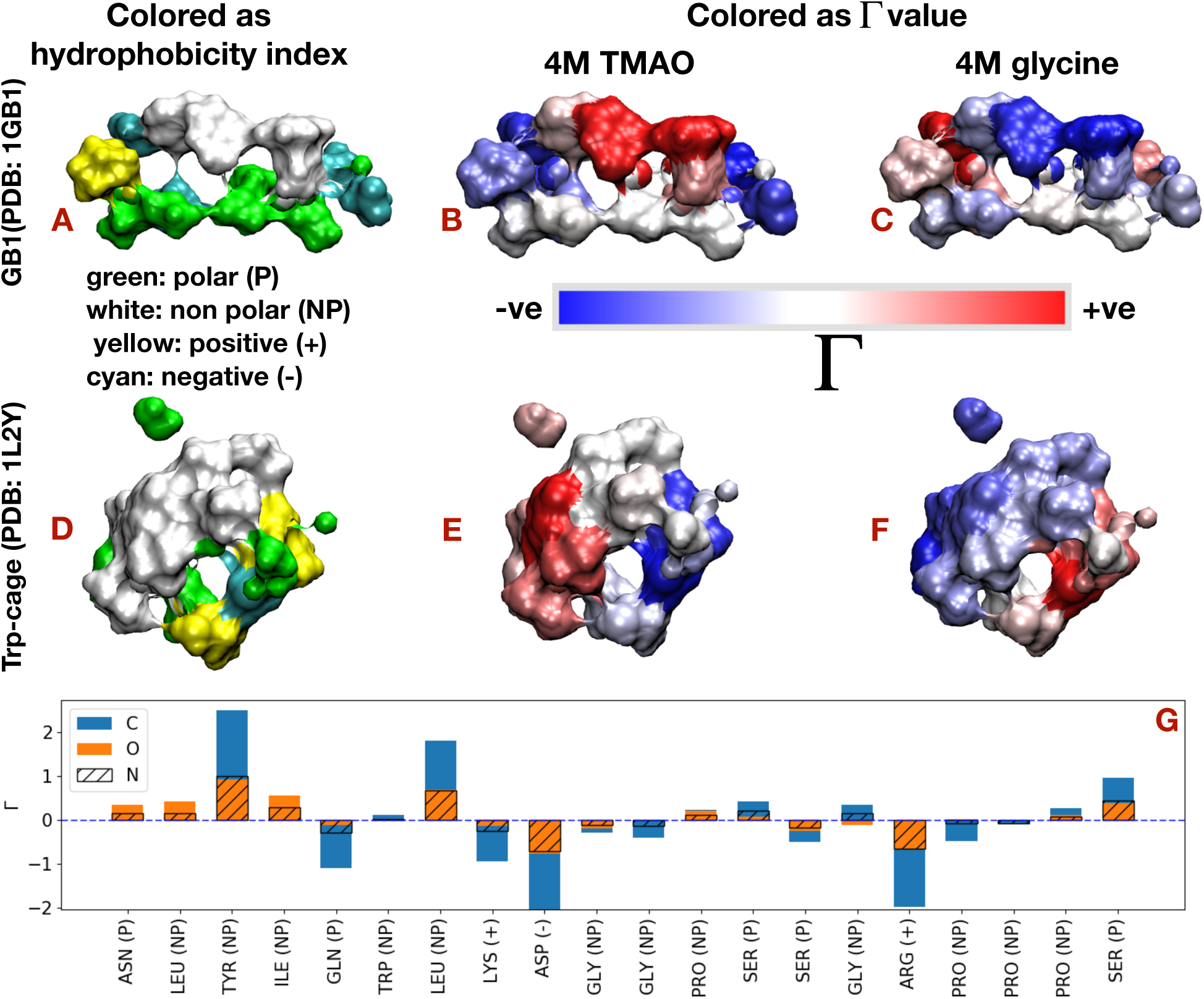
Surface map of side chain of constituent amino-acid residues of GB1 (A) and Trp-cage (D) according to the side chain hydrophobicity index. Similar map according to the preferential interaction (Γ) of side chain at *r* = 0.55 nm of constituent amino-acid residues of GB1 and Trp-cage in aqueous solution of TMAO (B and E) and in aqueous solution of glycine (C and F). (G) TMAO Γ values at *r* = 0.55 nm computed using its constituent atoms (methyl carbon (C), oxygen (O) and Nitrogen (N) atoms) with each amino-acid side chains of Trp-cage.

### Conformation-dependent preferential interaction drives osmolyte-induced protein stabilisation

The present reports of ambivalent preferential interaction of TMAO with multiple mini proteins (preferential binding with Trp-cage vs preferential exclusion in GB1) strike apparently contrasting view with the popular osmophobic model. However, here we show that this apparent conflict is reconciled when we appeal to the preferential interaction theory proposed by Tanford and Wyman.^19,20^ Based on this theory, the effect of preferential binding on a conformational equilibrium between the folded and the Unfolded configurations C ⇄ E (with an equilibrium constant *K*) is usually interpreted in terms of the thermodynamic analysis put forward by Wyman and Tanford, ^19,20^ which leads to

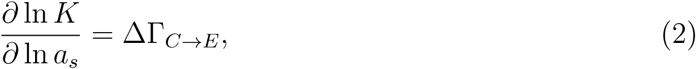

where *a*_*s*_ is the activity of the cosolute in the binary solution and ΔΓ_*C→E*_ is defined by (Γ^*E*^ − Γ^*C*^). According to Eq. 2, an increase in the concentration of the cosolute would lead to the macromolecular unfolding if ΔΓ > 0, and in contrast would favor the folded state over the unfolded one if ΔΓ < 0.

In summary, this theory states that it is not the absolute value of preferential interaction coefficient (Γ), rather the *difference* in the value of preferential interaction of osmolyte with the unfolded ensemble, relative to that with folded ensemble of the protein (ΔΓ = Γ_*unfolded*_- Γ_*folded*_), is directly related to the free energy of folding. Hence, a negative ΔΓ value free energetically drives the folding process. Towards this end, we individually compute the preferential interaction profiles of TMAO with both folded and unfolded conformations of GB1 and Trp-cage (Figure 5A and B). We find that TMAO depletes from both folded and unfolded ensembles of GB1, albeit more from the unfolded ensemble. On the other hand, while the same osmolyte shows preferential binding with *both* folded and unfolded ensembles of Trp-cage for a wide range of protein-osmolyte separation, the extent of preferential binding of TMAO is higher with folded ensemble of Trp-cage than that with its unfolded ensemble, thereby leading to ΔΓ < 0 in both cases, irrespective of preferential binding or exclusion. On the other hand, a similar analysis of preferential interaction of glycine with both the proteins suggests its preferential exclusion from both unfolded and folded ensembles (Figure 5C and D). However, in both cases, we find that ΔΓ < 0, (i.e. more preferential exclusion from unfolded ensembles) which leads to favorable free energy of the folding process. Taken together, we find that it is not the qualitative picture of binding versus exclusion, rather the overall conformation dependence in the extent of preferential interaction of TMAO and glycine with folded and unfolded conformation, which drives the folding process.

**Figure 5:**
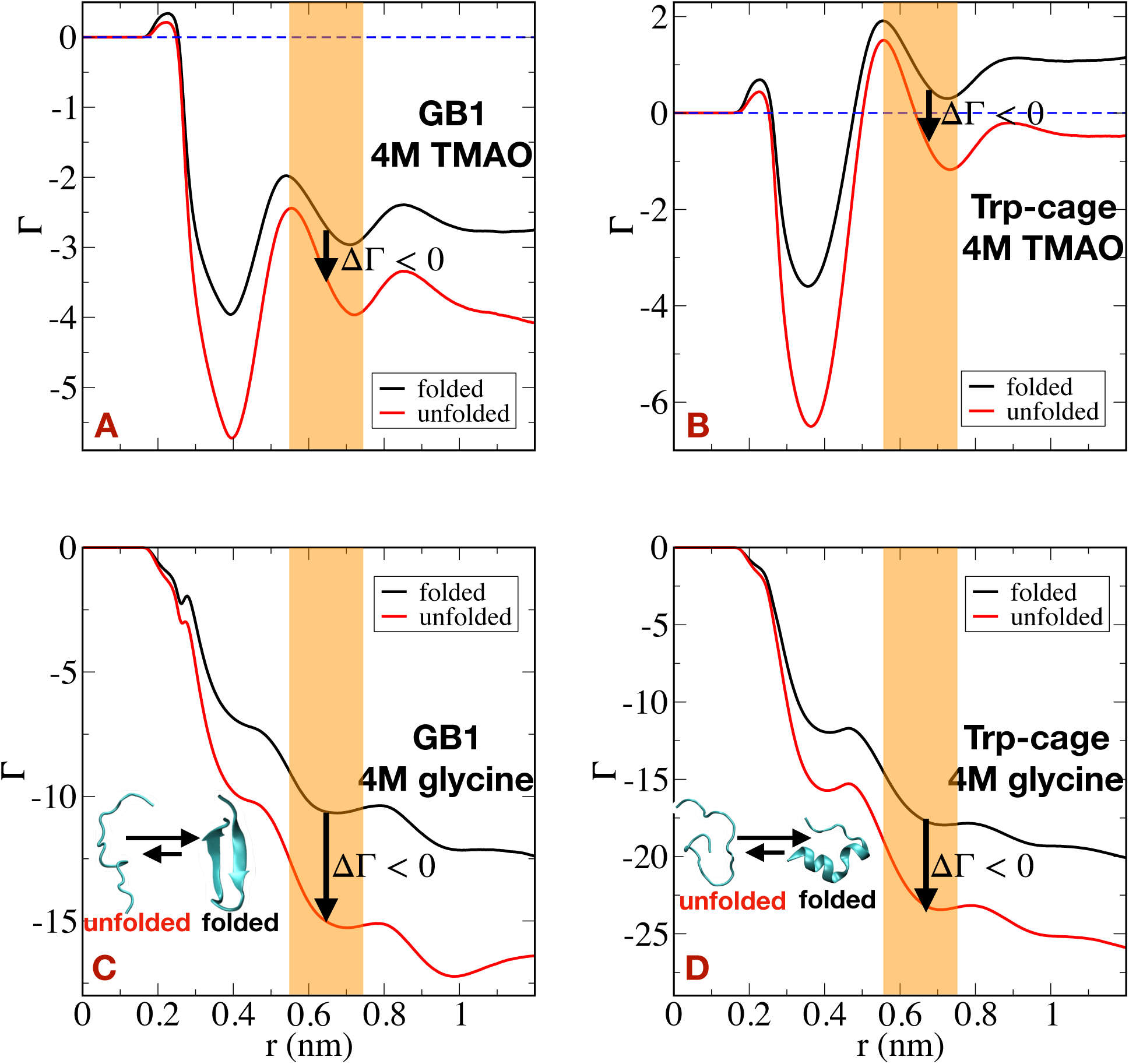
Preferential interaction coefficient (Γ) of TMAO (A and B) and glycine (C and D) with folded and unfolded conformations of GB1 and Trp-cage as a function of distance (*r*) from the protein surfaces.

### Towards establishing a common osmolyte-interaction motif across mini proteins

From the previous analysis, we inferred that the presence of methyl groups in TMAO, otherwise absent in glycine, encourages favorable hydrophobic interaction with the non polar side chains of GB1 and Trp-cage. Most importantly, we find that the presence of an extended hydrophobic patch in Trp-cage mini protein shifts the delicate balance in favor of net overall binding with the osmolytes. This led us to explore if the similar theme of osmolyte-protein interaction, especially the preferential binding of TMAO with protein, as observed in the case of Trp-cage, is prevalent in other archetypal mini proteins. Towards this end, we chose a set of well-known mini proteins (Figure 1C-F), namely Trp-zipper (PDB: 1LE1),^37^ Chignolin (PDB: 1UAO),^38^ Villin Headpiece (PDB: 1VII)^39^ and *βαβ* (PDB: 2KI0)^40^ for investigation of the preferential interaction of their folded state with TMAO and glycine. We simulate each of these proteins’ folded state in the aqueous solution of 4M glycine and TMAO and compute the respective preferential interaction profiles of these osmolytes with each of the proteins. Figure 6A-B compares the distance-dependent preferential interaction profiles of glycine and TMAO with a set of mini proteins. We find that, for all mini proteins under investigation here, Γ of glycine is negative for a full range of protein-osmolyte distance (Figure 6A), re-affirming glycine’s generic preferential exclusion from all proteins. However, on the other hand, the qualitative nature of TMAO’s preferential interaction varies with mini proteins (Figure 6B). Specifically, we find that while TMAO displays a clear trend of preferential exclusion from *βαβ*, consistent with the previous observation in GB1, it preferentially binds with Trp-zipper similar to that observed in Trp-cage. Villin Headpiece and Chignolin display somewhat intermediate signatures of weaker binding with TMAO, while mostly excluding from the protein. Mapping the residue-wise preferential interaction coefficient values of TMAO with the hydrophobicity index of constituent amino acids of each of the mini proteins recovers a common motif across all the mini proteins: hydrophobic amino acids display favorable binding with TMAO while polar side chains prefer to exclude TMAO from their surface and presence of extended hydrophobic domains leads to overall favorable interaction with TMAO (Figure 6G-J). Both Trp-zipper (Figure 6C) and Trp-cage (Figure 4D) possess an extended patch of hydrophobic residues, otherwise absent in GB1 (Figure 4A) and *βαβ* (Figure 6F), leading to apparent differences in their interaction with TMAO. The values of Γ of glycine are mostly zero to negative with most side chains (Figure 6K-N) due to the absence of methyl group in glycine.

**Figure 6:**
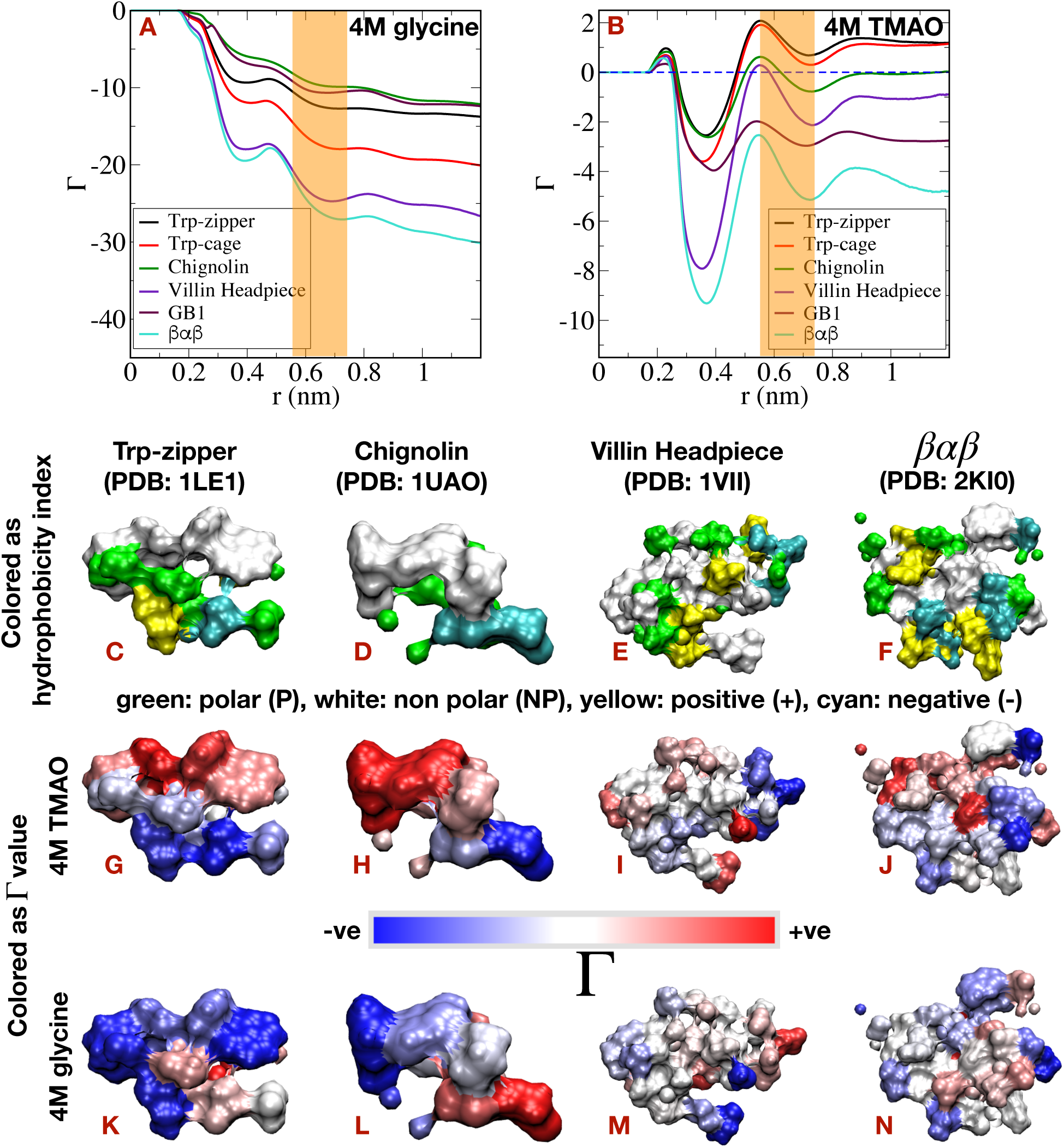
Preferential interaction coefficient (Γ) of glycine (A) and TMAO (B) with Trp-zipper, Trp-cage, Chignolin, Villin Headpiece, GB1 and *βαβ* as a function of distance (*r*) from the protein surfaces. Surface map of side chain of constituent amino-acid residues of Trp-zipper (C), Chignolin (D), Villin Headpiece (E) and *βαβ* (F) according to the side chain hydrophobicity index. Similar map according to the preferential interaction (Γ) of side chain at *r* = 0.55 nm of constituent amino-acid residues of those mini proteins in aqueous solution of TMAO (G-J) and in aqueous solution of glycine (K-N).

## Conclusion

In summary, we have investigated the role of the aqueous solution of glycine and TMAO on the folding equilibria of multiple prototypical mini proteins. We find that both the osmolytes (glycine and TMAO) stabilize the folding configurations of the proteins with respect to the neat eater. But glycine and TMAO behave differently in terms of preferential interaction to the surface of the folding configuration of mini proteins. Specifically, in the case of GB1, both glycine and TMAO exclude from the protein surface. But, glycine excludes from the surface of Trp-cage, while TMAO accumulates to the surface of Trp-cage, in contrast to the classical view of osmolyte-induced protein folding.

The origins of current anomaly from existing classical models can be traced to two salient features that were unraveled from the current work: first, we find that, while osmolytes always exclude from the backbone of the proteins, in congruence with the osmophobic model, it is the side chain of the proteins which dictates the overall preferential interaction behavior of the osmolytes. Detailed amino acids based preferential interaction calculation reveals that the presence of an extended patch of hydrophobic groups in Trp-cage leads to the overall accumulation of TMAO to the protein surface due to the hydrophobic interaction of the methyl group of TMAO and the hydrophobic side chain of the protein. On the other hand, these hydrophobic residues are dispersed among polar and charged residues in the case of GB1, leading to unfavorable interaction with TMAO. In the case of glycine, the overall preferential interaction coefficient is negative due to the absence of methyl group in glycine. We have also investigated this idea of favorable hydrophobic interaction with TMAO in the case of several mini proteins, namely Trp-zipper, Chignolin, Villin Headpiece and *βαβ*. We find that consistent preferential binding/exclusion of osmolytes to/from the protein surface depends on the topology of the surfaces of the folding configuration of the proteins. Secondly and most importantly, we show that the apparent ambivalence of osmolyte’s behavior towards protein and its implication in osmolytes’ stabilization role of protein can be unified by quantifying *conformation dependent* preferential interaction of osmolytes with the protein surface, via classic Wyman-Tanford theory.^19,20^ The current work convincingly demonstrates that irrespective of preferential binding or exclusion of osmolytes, it is the extent of preferential interaction of both the osmolytes which is more favorable with the folded ensemble of proteins than that with its unfolded ensemble, thereby shifting the overall folding equilibrium towards folded configurations of these mini proteins.

We note that there are precedent reports of preferential binding of TMAO to the protein surface. In particular, previous investigation by Su and coworkers^17^ has also reported possibility of preferential binding of TMAO to the protein surface. But their suggested mechanistic hypothesis of protein backbone getting stabilized by TMAO interaction via charge-charge interaction, at the cost of destabilization of hydrophobic interaction, is in contrast to earlier investigations on interaction of TMAO with hydrophobic proteins,^16^ which has suggested preferential binding of TMAO to the protein surface via stabilization of hydrophobic interaction in a surfactant-like mechanism. However, none of these previous works has quantified the crucial conformation-dependent osmolyte preferential interactions. Rather it was the average conformation of the protein which was analyzed for preferential interaction. One of the possible cause for lack of any conformational dependent preferential interaction analysis in prior works might lie in the choice of protein’s radius of gyration as their sole metric for analyzing protein folding. However, as observed in the present work and earlier works,^32–34^ the major conformational changes in the proteins studied are captured mostly along fraction of native contacts, with the radius of gyration playing a secondary work. The current work’s ability to investigate osmolytes’ interaction with both the folded and unfolded ensembles of proteins in equal footing mainly stems from the quantitative elucidation of full protein folding landscape in presence/absence of osmolytes via extensive usage of enhanced sampling techniques, judicious choice of multiple folding collective variables, in combination with Markov-based statistical model.

The observation of heterogenous mechanisms of protein-stabilizing osmolytes, as in the current work, raises fresh questions on how it influences the mechanistic action of the mixtures of these osmolytes with urea. The recent investigations have shed interesting insights into the non-additive roles of TMAO-urea mixtures on the conformational landscape of proteins^41,42^ and hydrophobic interactions. ^25^ It will be interesting to see how the proposed conformation dependence of osmolytes’ individual preferential interaction with protein plays out in the case of mixtures. Moreover, the current work’s demonstration that protein-stabilizing osmolyte may have ambivalent interactions with the proteins, can have possible implication in their mechanistic action in modulating protein-protein interactions.^29,43–45^ Recent experiments in these directions have shown that osmolyte and non-osmolytes may have a distinct role in perturbing charge-mediated inter-protein interactions^43^ and that zwitterionic osmolytes can resurrect the electrostatic interactions among charged surfaces, with low temperature playing an important role.^44,46,47^ Future investigation in the possible molecular mechanism of osmolyte-induced protein-protein interaction and its possible modulation of liquid-liquid protein phase separation^48^ can provide interesting insights.

## Materials and methods

In this work, we have individually investigated the free energetics of folding process of two prototypical mini proteins (Figure 1A-B), namely GB1 *β*-hairpin (PDB: 1GB1, residues 41−56; we call it GB1)^30^ and Trp-cage (PDB: 1L2Y)^31^ in neat water, 4 M aqueous solution of glycine and 4 M aqueous solution of trimethylamine N-oxide (TMAO). Both the proteins were capped with an acetyl group on the N-terminal and methyl amide on the C-terminal. The systems were solvated with TIP3P^49^ water molecules and corresponding osmolytes (glycine or TMAO) and then charge neutralized by counter-ions (Na^+^ or Cl^−^). The systems were first energy minimized and eventually equilibrated for 0.1 ns in NVT ensemble at an equilibrium temperature of 303 K using the Nose-Hoover thermostat^50,51^ with time constant 1 ps and then equilibrated for 2 ns in NPT ensemble at equilibrium temperature of 303 K using the Nose-Hoover thermostat with a time constant 1 ps and the equilibrium pressure of 1 bar using the Berendsen barostat^52^ with time constant 1.0 ps and then again equilibrated for 10 ns in NPT ensemble at equilibrium temperature of 303 K using the Nose-Hoover thermostat with time constant 1 ps and the equilibrium pressure of 1 bar using the Parrinello-Rahaman barostat^53^ with time constant 1.0 ps. The particle-mesh Ewald (PME)^54^ method (grid spacing = 0.12 nm) was used for long-range electrostatics. LINCS^55^ method and SETTLE^56^ algorithm were used to constrain the bonds associated with Hydrogen and the bonds and angle of water molecules respectively. GROMACS 2018^57^ software was used to perform all the molecular dynamics (MD) simulations.

The previously equilibrated systems were used to perform replica exchange molecular dynamics simulation (REMD) ^58^ with a total of 56 replicas in the temperature range of 280 − 540 K. Each of the replicas was initially equilibrated for 10 ns in NVT ensemble at an equilibrium temperature of 303 K using the Nose-Hoover thermostat with a time constant 1 ps and subsequently the simulations were carried out for 400 ns with a replica exchange interval of 10 ps. An average exchange probability of 0.25 was obtained.

The REMD conformations at 303 K were subsequently divided into 200 clusters using the backbone RMSD based k-means clustering algorithm. ^59^ Further 200 independent MD simulations of 100 ns each were carried out starting from the different initial configurations selected randomly from individual clusters.

The 200 MD short simulation trajectories for both the proteins GB1 and Trp-cage eventually used separately to build a Markov state model (MSM) ^60^ for statistically mapping the complete protein folding process using Pyemma software.^61,62^ The stationary populations of the discrete microstates, calculated from MSM, were subsequently used to reweigh the free energy surfaces derived from these short trajectories.

The radius of gyration (*r*_*g*_) was calculated using GROMACS software analysis tools and the fraction of native contacts (*NC*) was calculated using PLUMED^63^ software with a cutoff of 0.65 nm.

Additional 10 independent short MD simulations of 50 ns were carried out for unfolded (large *r*_*g*_ and small *NC*) and folded states (small *r*_*g*_ and large *NC*) separately with the 10 different initial configurations for each case (unfolded or folded states) selected from the unfolded and folded minima of the free energy surfaces by restraining the configurations of the proteins to their initial states. The last 40 ns of each trajectory were used to calculate the preferential interaction coefficients (defined in Results and Discussion section). The shaded regions in the distance-dependent preferential interaction coefficients (see main text) were considered based on the pair correlation function between osmolyte-protein (the region between the first maximum and the second minimum of pair correlation function).

We have also investigated the trend of preferential interactions of osmolytes with a set of other mini proteins (Figure 1C-F), namely Trp-zipper (PDB: 1LE1),^37^ Chignolin (PDB: 1UAO),^38^ Villin Headpiece (PDB: 1VII)^39^ and *βαβ* (PDB: 2KI0).^40^ Each of these mini proteins’ crystal structures were solvated, capped with an acetyl group on the N-terminal and methyl amide on the C-terminal and were first equilibrated at an equilibrium temperature of 303 K and the equilibrium pressure of 1 bar using the same protocol using in case of GB1 or Trp-cage. 4 M aqueous solution of glycine and 4 M aqueous solution of TMAO were used for each mini proteins. Subsequently, 10 short independent MD simulations of 50 ns in NVT ensemble at equilibrium temperature of 303 K were carried out for each case by restraining the configurations of these mini proteins to their native folded states. Similarly, the last 40 ns of each trajectory were used to calculate the preferential interaction coefficients.

In the majority of the works, the force field developed by Shea and coworkers,^12,64^ was used for TMAO and CHARMM36^65^ force field was used for glycine, proteins and counterions. Additional simulations were carried out using Kast^66^ and Netz^67^ force fields of TMAO to calculate preferential interaction coefficients in the case of GB1 and Trp-cage. We also tested the robustness of result with Trp-cage by using a different protein force field AMBER99SB-ILDN.^36^ Details of the simulations (number of molecules) for all the mini proteins are given in Table 1.

**Table 1:** Details of the simulations performed in the manuscript for the mini proteins.

| Proteins | Number of TMAO/<br>glycine molecules | Number of water<br>molecules in presence<br>of TMAO/glycine | TMAO/glycine<br>concentration |
| --- | --- | --- | --- |
| **GB1** | 0<br>205 | 2130<br>1747/1933 | 0 M<br>4 M |
| **Trp-cage** | 0<br>180 | 1823<br>1487/1662 | 0 M<br>4 M |
| **Trp-zipper** | 180 | 1521/1692 | 4 M |
| **Chignolin** | 180 | 1554/1724 | 4 M |
| **Villin Headpiece** | 180 | 1415/1584 | 4 M |
| $\beta\alpha\beta$ | 180 | 1419/1582 | 4 M |

## Supplemental Material

Supporting figures illustrating the robustness of the results across different osmolyte and protein force fields.

## Acknowledgements

This work was supported by computing resources obtained from shared facility of TIFR Center for Interdisciplinary Sciences, India. JM would like to acknowledge research intramural research grants obtained from TIFR, DAE, India, Ramanujan Fellowship and Core research grant provided by the Department of Science and Technology (DST) of India (CRG/2019/001219).

---

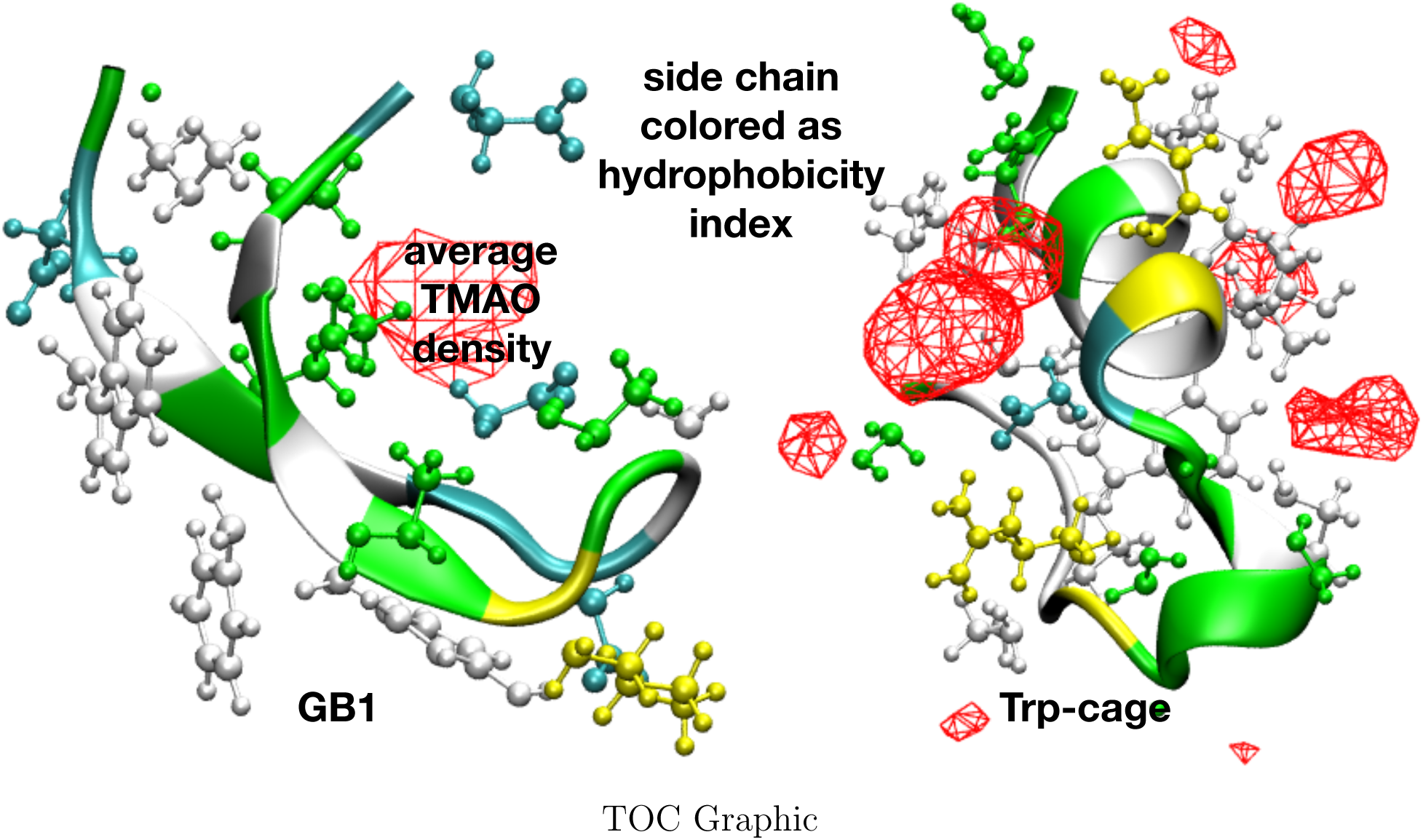

## Supporting information for “Unifying Heterogenous Mechanism of Protein-stabilising Osmolytes”

**Figure S1:**
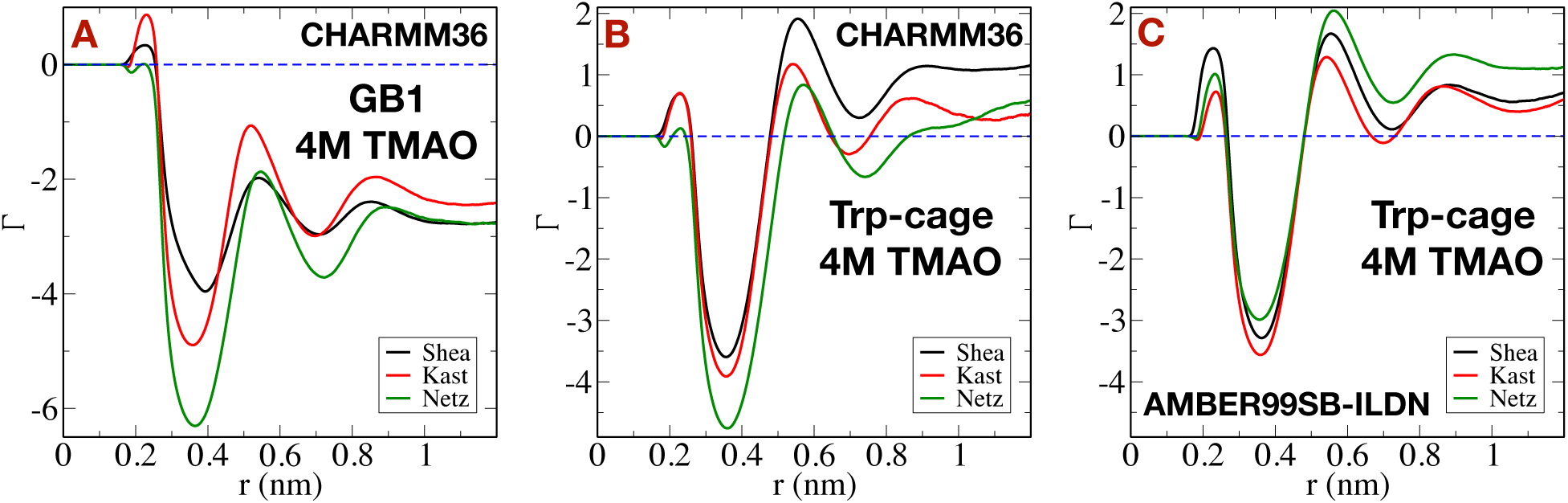
Preferential interaction coefficient (Γ) of TMAO with GB1 (A) and Trp-cage (B) as a function of distance (*r*) from the protein surfaces with the variation of TMAO force fields (namely “Shea”, “Kast” and “Netz”) with the protein force field CHARMM36. (C) Similar plot for Preferential interaction coefficient (Γ) of TMAO with Trp-cage with the protein force field AMBER99SB-ILDN

